# Estimating the distribution of fitness effects in aye-ayes (*Daubentonia madagascariensis*), accounting for population history as well as mutation and recombination rate heterogeneity

**DOI:** 10.1101/2025.01.02.631144

**Authors:** Vivak Soni, Cyril J. Versoza, Susanne P. Pfeifer, Jeffrey D. Jensen

## Abstract

The distribution of fitness effects (DFE) characterizes the range of selection coefficients from which new mutations are sampled, and thus holds a fundamentally important role in evolutionary genomics. To date, DFE inference in primates has been largely restricted to haplorrhines, with limited data availability leaving the other suborder of primates, strepsirrhines, largely under-explored. To advance our understanding of the population genetics of this important taxonomic group, we here map exonic divergence in aye-ayes (*Daubentonia madagascariensis*) – the only extant member of the Daubentoniidae family of the Strepsirrhini suborder. We further infer the DFE in this highly-endangered species, utilizing a recently published high-quality annotated reference genome, a well-supported model of demographic history, as well as both direct and indirect estimates of underlying mutation and recombination rates. The inferred distribution is generally characterized by a greater proportion of deleterious mutations relative to humans, providing evidence of a larger long-term effective population size. In addition however, both immune-related and sensory-related genes were found to be amongst the most rapidly evolving in the aye-aye genome.

## MAIN

The distribution of fitness effects (DFE) summarizes the range of selection coefficients from which new mutations are sampled. Consequently, characterizing the DFE holds a fundamentally important role in evolutionary genomics, as it quantifies the fraction of neutrally evolving genomic mutations, provides insights into the expected relative frequencies of purifying relative to positive selection, and informs the expected effects of selection at linked sites, to name but a few implications (see reviews of Eyre-Walker and Keightley 2007; Keightley and Eyre-Walker 2010; Bank et al. 2014a). Moreover, given that the vast majority of fitness-impacting mutations are deleterious, the constant elimination of these variants via purifying selection and the associated background selection (BGS) effects (Charlesworth et al. 1993) represent constantly operating processes shaping levels and patterns of genomic variation in and around functional regions. As such, an accurate characterization of these effects is critical for the construction of any evolutionary baseline model for a given species (Comeron 2014, 2017; Johri et al. 2022a; Howell et al. 2023; Terbot et al. 2023; Soni et al. 2023; Soni and Jensen 2024), and, because these effects may differ strongly depending on the relative proportion of weakly relative to strongly deleterious mutations, the DFE shape again emerges as a fundamental component for any evolutionary modeling or inference (Charlesworth et al. 1993; Hudson and Kaplan 1994; Charlesworth et al. 1995; Ewing and Jensen 2014, 2016; Johri et al. 2020).

Generally speaking, there are two classes of DFE inference, one applicable to lab-tractable organisms that may be experimentally evolved, and one applicable to natural population analysis. The former includes mutation accumulation experiments – in which a population of organisms can be maintained often in replicate, sampled at regular intervals, and the fitness effects of newly arising mutations characterized with respect to, for example, the wild-type state (e.g., Lenski et al. 1991; Barrick and Lenski 2013; Desai 2013; Böndel et al. 2019; Morales-Arce et al. 2022; Crombie et al. 2024). This class also includes mutagenesis experiments – in which hundreds or thousands of individuals can be maintained that carry one or very few mutations, and their fitness assessed by, for example, relative growth rates (e.g., Hietpas et al. 2011; Jacquier et al. 2013; Bank et al. 2014b; Fowler and Fields 2014; Matuszewski et al. 2015). Both methods represent powerful DFE inference approaches for the organisms in which they can be applied (e.g., *Saccharomyces cerevisiae, Caenorhabditis elegans, Chlamydomonas reinhardtii*), with the caveat being that they provide DFE inference only within the context of a lab-grown environment.

With regards to natural population analysis, which will be our focus here, there are generally approaches utilizing divergence data, polymorphism data, or a combination of both. Perhaps the most basic approach utilized to infer aspects of the DFE relies on comparisons between non-synonymous and synonymous divergence. Assuming that synonymous sites are effectively neutral, and thus characterized by a substitution rate equal to their mutation rate (Kimura 1968), one may quantify the fraction of non-synonymous mutations that are deleterious (and thus characterized by reduced fixation probabilities relative to neutrality) by assessing the depression in non-synonymous divergence relative to the synonymous neutral standard (e.g., Eyre-Walker et al. 2002). Similarly, if advantageous mutations are present (characterized by increased fixation probabilities relative to neutrality), one may assess this fraction of the DFE via the acceleration of non-synonymous divergence relative to synonymous (e.g., Smith and Eyre-Walker 2002). These advances largely owed to the realization that a McDonald-Kreitman-style test (McDonald and Kreitman 1991) could be used to infer proportions of adaptive substitutions (Charlesworth 1994). Synonymous and non-synonymous mutations aside, one may similarly utilize this divergence-based logic to assess selective constraints acting in different genomic regions (e.g., coding relative to intronic relative to intergenic; Andolfatto 2005).

When incorporating polymorphism data into DFE inference, one initial challenge is the need to incorporate the demographic history of the population into the inference procedure, given that this history may also act to shape levels and patterns of variation and thus may potentially result in mis-inference if unaccounted for (see review of Johri et al. 2022b). One of the first advances in this regard utilized the site frequency spectrum (SFS) at putatively neutral synonymous or non-coding sites in order to infer a population history, and then conditioned on that history to infer the DFE at putatively functional non-synonymous sites (Williamson et al. 2005; Keightley and Eyre-Walker 2007). Such step-wise approaches yielded some of the first polymorphism-based DFE estimates for a variety of organisms (Eyre-Walker and Keightley 2007; Boyko et al. 2008; Eyre-Walker and Keightley 2009; Schneider et al. 2011). A related category of methods also arose for utilizing time-sampled polymorphism data in order to infer individual mutational effects based on observed allele frequency changes – as may be applicable to ecological datasets or ancient DNA sampling – with the stochastic effects of genetic drift associated with the given population history being incorporated by estimating an effective population size based on the variance observed in neutral allele frequencies (e.g., Malaspinas et al. 2012; Foll et al. 2015; Ferrer-Admetlla et al. 2016; and see review of Malaspinas 2016). However, in addition to generally being limited to relatively simple population-size change models (though more complex models have been developed; e.g., Ma et al. 2013; Kim et al. 2017), these initial single- and multi-timepoint approaches also assume independence amongst sites, and thus neglect any role of background selection or other forms of genetic hitchhiking in further shaping levels of polymorphism (see reviews of Charlesworth and Jensen 2021, 2022).

In order to address these polymorphism-based challenges, simultaneous inference approaches have recently been developed. Though accounting for the effects of selection on linked sites within an analytical framework remains challenging, Cvijovic et al. (2018) obtained expressions for the SFS at sites experiencing BGS in a constant size population, and Friedlander and Steinrücken (2022) described a numerical framework to obtain expected SFS and linkage disequilibrium (LD) patterns around a selected region with changing population size. In order to allow for more complex models, progress has also been made using approximate Bayesian computation (ABC) approaches with forward simulations, in order to model both complex population histories and flexible DFE shapes, whilst accounting for the resulting effects of selection on linked sites. For example, Johri et al. (2020) developed a joint ABC approach estimating the DFE densities of neutral, weakly deleterious, moderately deleterious, and strongly deleterious mutations, together with a history of population size change, utilizing aspects of the SFS, LD, and divergence as summary statistics. Notably, the exclusion of BGS effects in previous methods was found to result in an under-estimation of weakly deleterious mutations and an over-estimation of population growth, a bias which is corrected within this ABC framework (Johri et al. 2021). Subsequent work has also demonstrated the potential mis-inference that may arise by neglecting underlying heterogeneity in rates of both mutation and recombination (Soni et al. 2024a). Taken together, this literature thus emphasizes the importance of incorporating population history, the effects of selection at linked sites, and mutation and recombination rate maps / uncertainties when performing DFE inference.

In primates specifically, these various divergence- and polymorphism-based approaches have been employed widely, with humans being the best studied in this regard. For example, Keightley and Eyre-Walker (2007) fit a gamma-distributed DFE utilizing a gene set associated with severe disease or inflammatory response, and estimated a large proportion (∼40%) of strongly deleterious mutations and a relatively low (∼20%) proportion of effectively neutral mutations. Huber et al. (2017) utilized a wider selection of genes, resulting in a DFE skewed towards effectively neutral mutations (∼50%; similar to the estimate of Johri et al. 2023 utilizing a different subset of genes), and a smaller proportion of strongly deleterious mutations (∼20%). Thus, these differences may well simply and accurately reflect true DFE differences in the underlying gene sets evaluated. Similar inference has also been performed across the great apes (e.g., Castellano et al. 2019; Tataru and Bataillon 2020), and considerations have been extended to general regulatory regions as well (e.g., Simkin et al. 2014; Anderson et al. 2020; Kuderna et al. 2024).

Notably however, owing largely to data availability, these estimates have been performed primarily in haplorrhines, with the other suborder of primates, strepsirrhines, being largely unexplored. Yet, a number of recent advances have uniquely enabled investigation in this neglected space of the primate clade. Firstly, Versoza & Pfeifer (2024) have recently provided an annotated chromosome-level genome assembly for aye-ayes (*Daubentonia madagascariensis*), thereby allowing for the essential demarcation of functional and non-function genomic regions needed for performing DFE inference. Secondly, recent work has also generated high-quality direct mutation and recombination rates for aye-ayes from multi-generation pedigree data (Versoza et al. 2024a,b; Versoza, Lloret-Villas, et al. 2024) – as well as indirect fine-scale estimates based on autosomal patterns of LD and neutral divergence (Soni, Versoza, et al. 2024). Finally, utilizing high-coverage whole-genome data from unrelated individuals, Terbot et al. (2024) recently estimated a well-fitting population history for aye-ayes (and see Soni et al. 2024b) – which described a severe and ancient population size decline likely associated with the human colonization of Madagascar, as well as a more recent decline likely associated with habitat destruction – thereby providing the needed accounting of the role of population history in shaping observed SFS across the genome.

### Interpreting exonic divergence

Building upon these recent results, we replaced the original aye-aye genome in the 447-way mammalian multiple species alignment (Zoonomia Consortium 2020) that includes hundreds of closely related primate species (Kuderna et al. 2023) with the high-quality aye-aye reference genome of Versoza and Pfeifer (2024) to quantify fine-scale exonic divergence in the species. Figure 1 summarizes observed exonic divergence on the aye-aye branch, with the maximum neutral divergence for 1kb and 1Mb windows (Soni, Versoza, et al. 2024) provided for orientation. Although a considerable number of exons were characterized by rates of fixation greater than the maximum neutral divergence observed in 1Mb windows, no exons were found to be in excess of the neutral rate observed in 1kb windows – the more appropriate comparison given that the mean exonic length is <1kb. Thus, exonic divergence was observed to be lower than the maximum neutral divergence in aye-ayes without exception, as expected from the dominant action of purifying selection in functional regions (Charlesworth et al. 1993).

**Figure 1:**
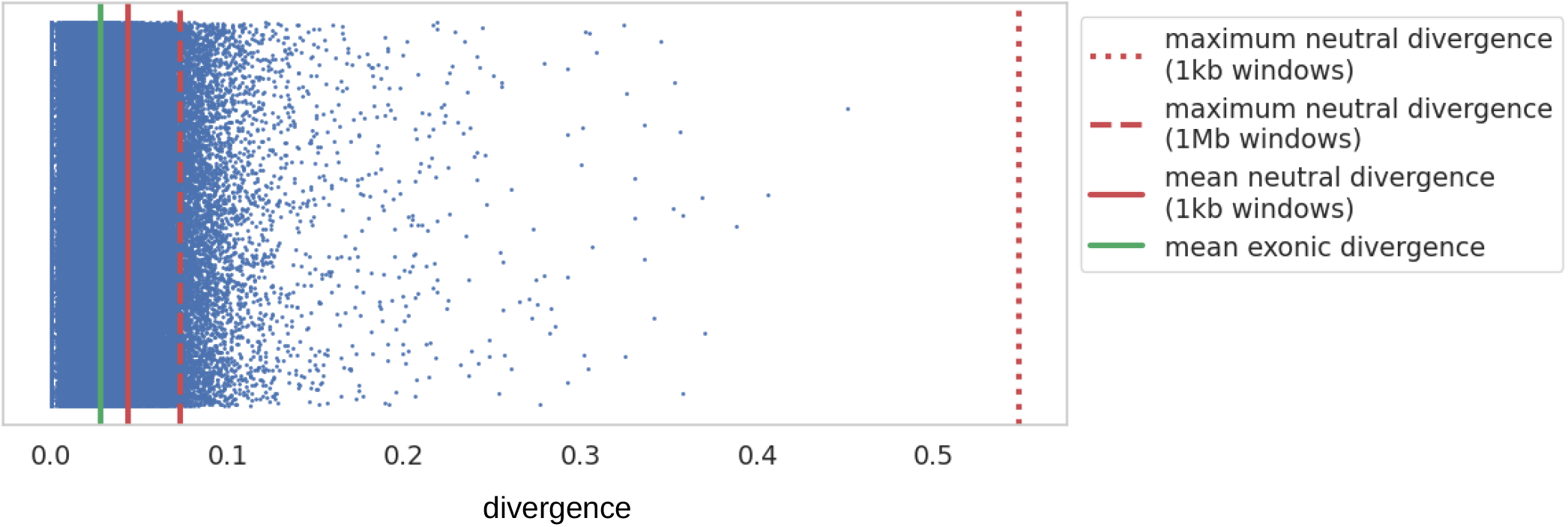
Exonic divergence scatter plot with maximum neutral divergence values marked for windows of size 1Mb (red dashed line) and 1kb (red dotted line), as well as the mean neutral divergence for 1kb windows (red solid line), as calculated from Soni, Versoza, et al. (2024). Exonic divergence was calculated by replacing the aye-aye in the 447-way mammalian multiple species alignment (Zoonomia Consortium 2020; Kuderna et al. 2023) with the high-quality aye-aye reference genome of Versoza and Pfeifer (2024), via the Cactus alignment software (Armstrong et al. 2020), and extracting divergent sites along the aye-aye branch using the HAL software package (Hickey et al. 2013). Exonic divergence was calculated by dividing the number of divergent sites in each exon by the total accessible exonic length, and only exons with a minimum of 100bp of accessible sites were included. Each dot represents an autosomal exon, and the mean exonic divergence is plotted (green solid line).

However, given that even recurrent positive selection is still expected to be rare relative to purifying selection, the absence of entire exons evolving faster than neutrality does not itself eliminate the possibility of positive selection contributing to exonic divergence in subsets of genes. For example, distinct classes of exons were found to occupy the tails of the exonic divergence distribution, and were found to be in excess of the mean neutral fixation rate. Utilizing the genome annotations from the Versoza and Pfeifer (2024) reference genome to calculate the mean divergence per gene, we ran a gene functional analysis using g:Profiler (Kolberg et al. 2023) on all coding regions with a mean divergence greater than the 75^th^ percentile of neutral divergence (0.039694). Figure 2 provides the divergence distribution of all examined exons, compared with the distributions of the two fastest-evolving gene classes – those related to sensory and immune function.

**Figure 2:**
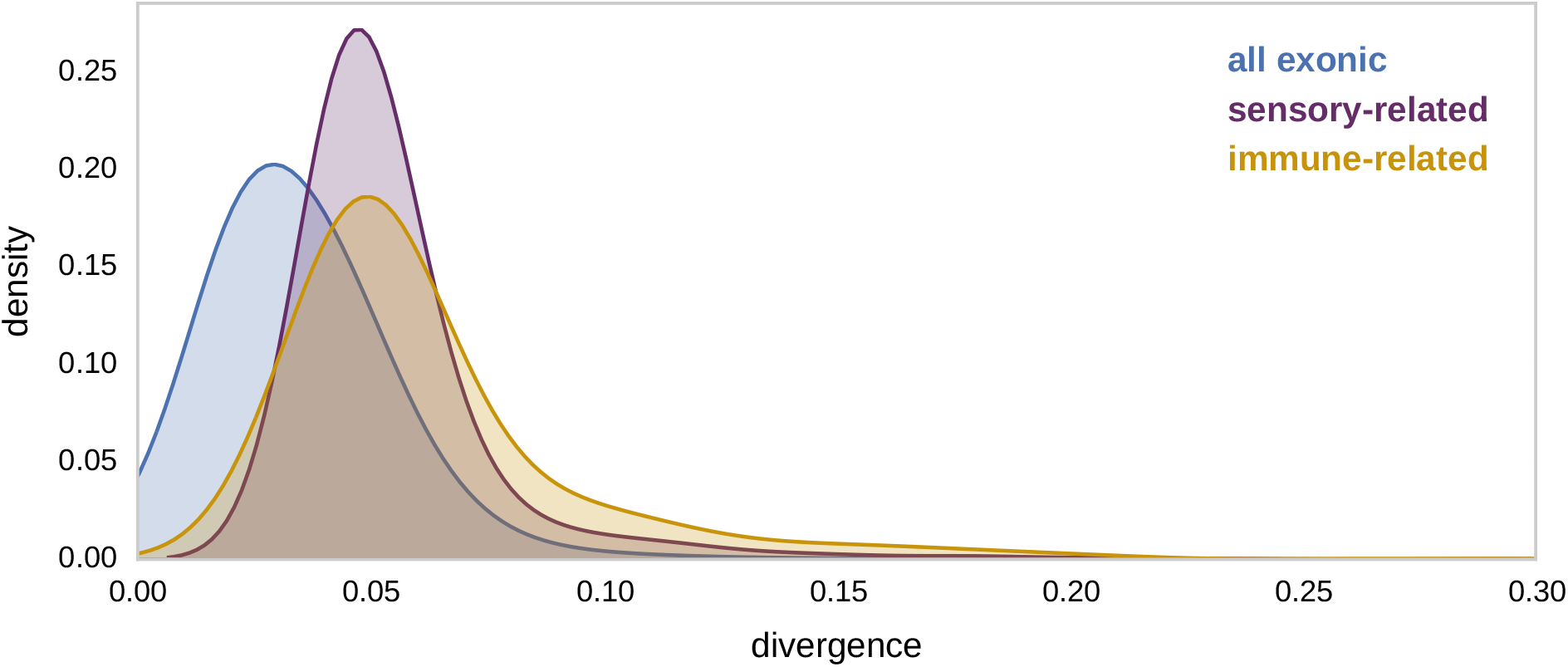
Density plots of exonic divergence in aye-ayes for all exons (blue), exons located in genes implicated in sensory-related functions (purple), and exons located in genes implicated in immune-related functions (gold). Utilizing the Versoza and Pfeifer (2024) genome annotations, the mean divergence per gene was calculated, and gene functional analysis was performed using g:Profiler (Kolberg et al. 2023) on the set of genes with a divergence value greater than the 75^th^ percentile of neutral divergence in aye-ayes (0.0397; Soni, Versoza, et al. 2024).

Immune-related genes have long been observed to be amongst the most rapidly evolving across vertebrates, as populations continually respond to challenges of pathogen exposure (e.g., George et al. 2011; Rausell and Telenti 2014), and our results remain consistent with this pattern. With regards to the sensory-related distribution, both the nocturnal activity patterns of aye-ayes (with the suggestion previously being made that dichromacy may enable aye-ayes to perceive color whilst foraging in moonlight conditions; Perry et al. 2007), together with evidence that aye-ayes may discriminate between individuals based on scent (Price and Feistner 1994) and use scent-marking to attract mates (Winn 1994), both suggest potentially significant roles for opsin- and olfactory-related genes throughout the evolutionary history of the species. Furthermore, Soni et al. (2024b) recently found that a number of sensory functional categories including G-protein coupled receptors and olfactory receptors had strong statistical support for being maintained by long-term balancing selection in aye-ayes – noting that diversity in these genes may increase the number of different odorant-binding sites (Lancet 1994) – further supporting these hypotheses.

### Utilizing patterns of exonic divergence to infer the DFE

Divergence is an informative summary statistic when inferring patterns of long-term selection, as the general features of the DFE are likely to remain relatively stable over deep evolutionary time. As such, we ran forward-in-time simulations in SLiM4.0.1 (Haller and Messer 2023) in order to fit observed empirical exonic divergence with a DFE shape consisting of neutral, weakly deleterious, moderately deleterious, and strongly deleterious mutational classes. In brief, we simulated a 54.9 million year divergence time of the aye-aye branch (Horvath et al. 2008), assuming a generation time of 5 years (Ross 2003; Louis et al. 2020) – both of which have additionally been recently supported by whole-genome neutral divergence patterns (Soni, Versoza, et al. 2024) – and utilized the estimated demographic model of Terbot et al. (2024) in order to characterize the recent history of the species. Using our multiple-species alignment to compute the number of divergent sites along the aye-aye branch, we were able to directly compare this empirical observation with the number of fixations accrued in our simulated population during the divergence phase.

As depicted in Figure 3, observed divergence was fit well by a DFE of new mutations characterized by a majority of nearly neutral variants, and a remaining even mix of moderately and strongly deleterious variants. For comparison, a recent estimate of the DFE from human populations (Johri et al. 2023) has also been included. As shown, humans were characterized by a higher density of neutral variants and a lower density of more strongly deleterious variants relative to aye-ayes, likely consistent with the smaller effective population size of the former.

**Figure 3:**
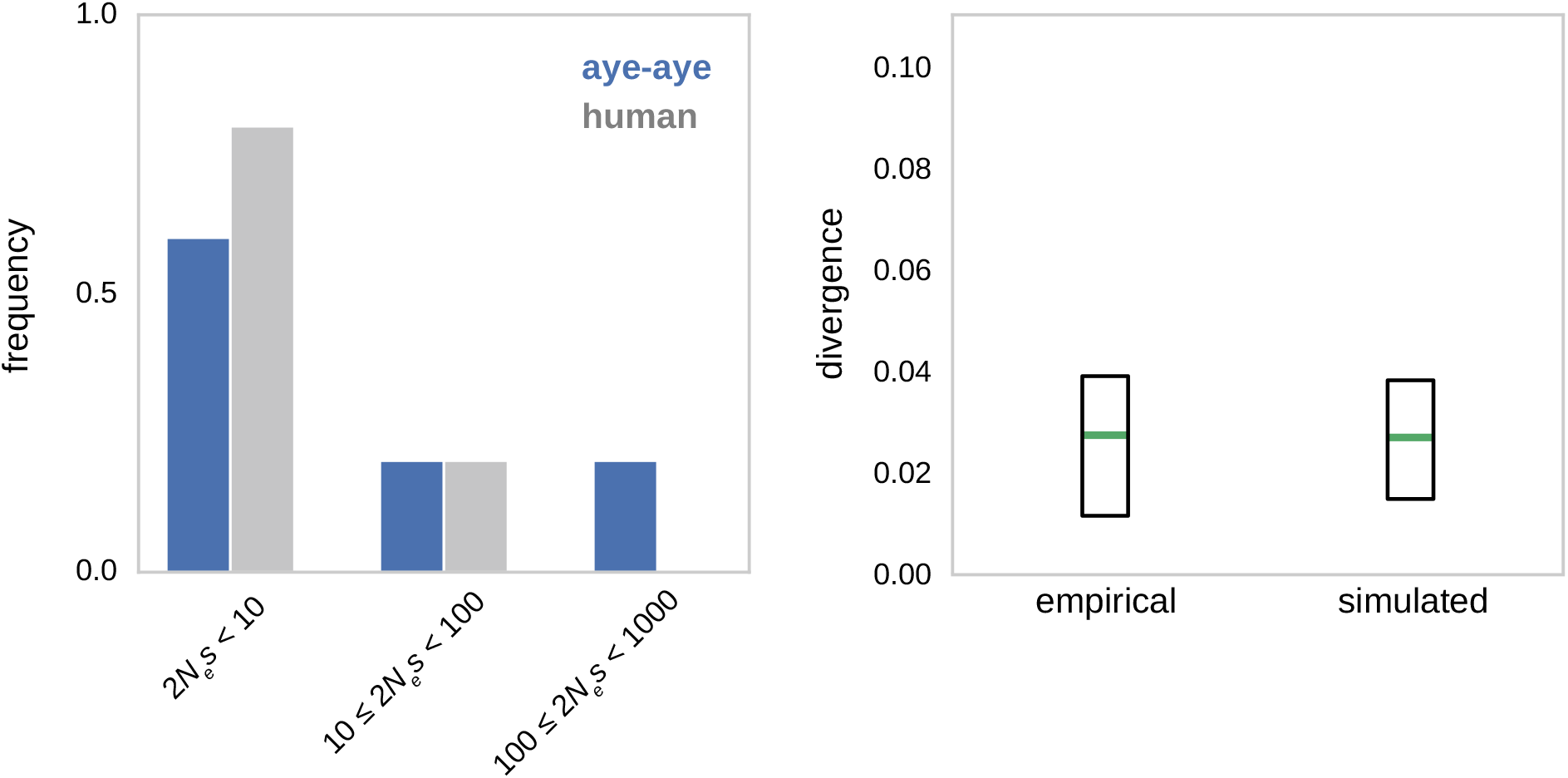
Comparison of the empirical and simulated divergence under the best-fitting DFE. Simulations were run in SLiM4.01 (Haller and Messer 2023) under the Terbot et al. (2024) aye-aye demographic model, assuming 54.9 million years since the branch split (Horvath et al. 2008) and a generation time of 5 years (Ross 2003; Louis et al. 2020). Simulations included a 10*N*_*ancestral*_ generation burn-in time prior to the demographic model (where *N*_ancestral_ is the initial population size and *s* the reduction in fitness of the mutant homozygote relative to wild type). For this example, the simulated exon length was 2,978bp (i.e., the mean empirical length of exons of size greater than 1kb). Mutation and recombination rate heterogeneity was modelled such that each simulated exon had a rate drawn from a normal distribution, with the mean rate across all 100 simulated exons equal to the mean pedigree estimated rates of 0.4e-8/bp/generation and 0.85cM/Mb for mutation and recombination, respectively (Versoza et al. 2024; Versoza, Lloret-Villas et al. 2024). Exonic mutations were drawn from a DFE comprised of four fixed classes (following Johri et al. 2020), denoted by *f*_*0*_ with 0 ≤ 2*N*_ancestral_ *s* < 1 (i.e., effectively neutral mutations), *f*_1_ with 1 ≤ 2*N*_ancestral_ *s* < 10 (i.e., weakly deleterious mutations), *f*_2_ with 10 ≤ 2*N*_ancestral_ *s* < 100 (i.e., moderately deleterious mutations), and *f*_3_ with 100 ≤ 2*N*_ancestral_ *s* (i.e., strongly deleterious mutations). **Left panel:** Best-fitting discrete DFE in ayes-ayes (blue), as compared to the Johri et al. (2023) DFE inferred for humans (grey). **Right panel:** Comparison of empirical and simulated divergence values for the best-fitting DFE. Green lines represent the mean value, whilst boxes represent 25 and 75 percentiles.

Notably however, this inference in aye-ayes assumes a mutation rate of 0.4e-8/site/generation, as was directly inferred from pedigree data (Versoza et al. 2024a). Yet, this pedigree inference was for the youngest parents in the study (9-11 years of age), and a strong parental age effect was observed. Namely, the oldest parents in the study (24-26 years of age) were characterized by a rate of 2.0e-8/site/generation, with an average rate across the pedigree of 1.1e-8/site/generation. Given that aye-ayes reach sexual maturity by 3 years of age (Winn 1994), reproduction in the wild likely occurs amongst individuals even younger than the youngest in the pedigree, and given support for the 0.4e-8/site/generation rate from recent indirect divergence-based inference (Soni, Versoza, et al. 2024), we believe this to be a reasonable estimate for our conversion here. However, if the true rate were to be even lower owing to parents being generally younger throughout the evolutionary history of the species, the inferred DFE would resultingly become more skewed towards nearly neutral variants and thus potentially more similar to the human estimate. It is also noteworthy that long-term mutation rates on the aye-aye branch higher than 0.4e-8/site/generation become very difficult to reconcile with the fossil-record (see Tavaré et al. 2002; Soni, Versoza, et al. 2024), consistent with the suggestion of generally lower rates in prosimians relative to other primates (see the reviews of Tran and Pfeifer 2018; Chintalapati and Moorjani 2020).

### Concluding summary

We have here characterized functional divergence in aye-ayes, finding, as expected, that exonic divergence is generally much reduced relative to neutral divergence. Yet, amongst exons, we also found evidence of an increased rate of divergence in genes related to immune and sensory-related functions, in agreement with previous work across vertebrates for the former, and in primates more specifically for the latter. Employing forward simulations to fit a DFE to observed exonic divergence, we found evidence of an increased proportion of newly arising deleterious variants in aye-ayes relative to humans, likely related to their larger estimated effective population size. These findings also generally support a relatively low mutation rate in aye-ayes compared to other primates, as has been proposed both from indirect neutral divergence as well as from direct pedigree-based inference.

## ACKNOWLEDGEMENTS

We would like to thank the Duke Lemur Center for providing the aye-aye samples used in this study, and members of the Jensen Lab and Pfeifer Lab for helpful discussion. Computations were performed on the Sol supercomputer at Arizona State University (Jennewein et al. 2023) and on the Open Science Grid, which is supported by the National Science Foundation and the U.S. Department of Energy’s Office of Science. This is Duke Lemur Center publication # XXXX.

